# A Methodology for Specific Disruption of Microtubules in Dendritic Spines

**DOI:** 10.1101/2024.03.04.583370

**Authors:** Elizabeth D. Holland, Hannah L. Miller, Matthew M. Millette, Russell J. Taylor, Gabrielle L. Drucker, Erik W. Dent

## Abstract

Dendritic spines, the mushroom-shaped extensions along dendritic shafts of excitatory neurons, are critical for synaptic function and are one of the first neuronal structures disrupted in neurodevelopmental and neurodegenerative diseases. Microtubule (MT) polymerization into dendritic spines is an activity-dependent process capable of affecting spine shape and function. Studies have shown that MT polymerization into spines occurs specifically in spines undergoing plastic changes. However, discerning the function of MT invasion of dendritic spines requires the specific inhibition of MT polymerization into spines, while leaving MT dynamics in the dendritic shaft, synaptically connected axons and associated glial cells intact. This is not possible with the unrestricted, bath application of pharmacological compounds. To specifically disrupt MT entry into spines we coupled a MT elimination domain (MTED) from the Efa6 protein to the actin filament-binding peptide LifeAct. LifeAct was chosen because actin filaments are highly concentrated in spines and are necessary for MT invasions. Temporally controlled expression of this LifeAct-MTED construct inhibits MT entry into dendritic spines, while preserving typical MT dynamics in the dendrite shaft. Expression of this construct will allow for the determination of the function of MT invasion of spines and more broadly, to discern how MT-actin interactions affect cellular processes.

**Significance Statement:**

- The LifeAct-MTED construct provides spatial and temporal control of microtubule depolymerization within individual cells
- Targeting this construct directly to spines allows for the specific inhibition of microtubule dynamics into dendritic spines, without affecting microtubule dynamics in the dendritic shaft
- Implementation of the construct will allow for the testing of microtubule contributions to specific biological processes, without global disruption of microtubule dynamics

## Introduction

Microtubule (MT) dynamic instability (Mitchison and Kirschner, 1984; Desai and Mitchison, 1997), the stochastic polymerization and depolymerization of MT polymers, occurs in all cell types, including mammalian neurons. However, as developing neurons mature into highly polarized cells containing elaborate dendrites and a single axon, it has been thought that MTs become stabilized to maintain neuronal architecture. With the discovery and fluorescent labeling of end-binding proteins that track along the tips of polymerizing MTs (Stepanova *et al*., 2003), it became apparent that a portion of cellular MTs continue to undergo polymerization and depolymerization for the life of the neuron (Hu *et al*., 2008; Yau *et al*., 2016). To discover the function of MT dynamics in neurons, researchers have used chemical compounds to inhibit MT dynamicity. At nanomolar concentrations, drugs such as paclitaxel (taxol), which stabilizes MTs, (Schiff and Horwitz, 1980) and nocodazole, which depolymerizes MTs, (Mareel and De Brabander, 1978), both inhibit MT dynamicity, resulting in little polymerization and depolymerization (Vasquez *et al*., 1997; Mikhailov and Gundersen, 1998; Yvon *et al*., 1999). Inhibiting MT dynamicity in this way has allowed for the discovery that MT dynamics are essential for many processes in developing neurons, including axon branching (Dent and Kalil, 2001).

More recently, several groups have shown that MTs have robust dynamics in mature cortical, hippocampal and cerebellar neurons, especially in dendrites (Hu *et al*., 2008; Jaworski *et al*., 2009; Wagner *et al*., 2011). These dynamic MTs polymerize throughout the dendritic arbor and can even enter the micron-sized, mushroom-shaped protrusions along dendrites called dendritic spines (Gu *et al*., 2008; Hu *et al*., 2008; Jaworski *et al*., 2009). In excitatory neurons, dendritic spines are the primary site of synaptic contact with presynaptic axons. Dendritic spines are not static structures but undergo morphological and molecular plasticity. They can undergo long-term potentiation (LTP) and enlarge or long-term depression (LTD) and shrink, depending on presynaptic activity (Tada and Sheng, 2006). Interestingly, MT entries into dendritic spines are dependent on neuronal activity (Gu *et al*., 2008; Hu *et al*., 2008; Jaworski *et al*., 2009) and result in long-term spine enlargement (Merriam *et al*., 2011). Indeed, MT invasion of spines increases during LTP and decreases during LTD (Kapitein *et al*., 2011; Merriam *et al*., 2013), showing a direct correlation with neuronal activity.

However, correlation does not indicate causation. To determine the function of MT invasion of dendritic spines requires inhibiting their entry into dendritic spines, while leaving the dynamics of MTs in the dendritic shaft, presynaptic axon and associated glial cells intact. Unfortunately, the primary methodology for inhibiting MT polymerization is the bath application of compounds such as paclitaxel and nocodazole. Addition of nanomolar concentrations of these drugs inhibits MT invasions of spines and results in decreased LTP (Jaworski *et al*., 2009), abolition of spine growth upon MT entry (Merriam *et al*., 2011), blockage of BDNF-dependent increase in PSD95 in spines (Hu *et al*., 2011) and inhibition of transport of a motor/cargo pair (KIF1A/synaptotagmin-IV) into dendritic spines (McVicker *et al*., 2016). Nevertheless, all these studies used bath applications of nocodazole or paclitaxel, which results in the inhibition of MT dynamics throughout the culture or slice preparation. Even if these pharmacological applications were localized to specific dendrites or spines, they would still affect the MTs present in the presynaptic axon. Therefore, a more spatially restrictive inhibition of MT dynamics is needed to definitively determine the function of MT invasion of dendritic spines.

Recently, others have designed optogenetic tools to depolymerize MTs, including photostatins (Borowiak *et al*., 2015), a construct based on the depolymerizing activity of kinesin 13 (Lu *et al*., 2020) or the MT severing enzymes spastin (Liu *et al*., 2022) or katanin (Meiring *et al*., 2022). Although these are useful tools to study the function of MTs, they do not lend themselves to the study of MT invasion of dendritic spines because MT invasions are infrequent and transient (Hu *et al*., 2008; Jaworski *et al*., 2009). It is also not possible to predict with any certainty which spine along a dendritic arbor will be invaded.

Thus, we developed a construct that targets and is maintained in all dendritic spines. This construct contains a potent MT elimination domain (MTED) peptide, derived from the Arf guanine nucleotide exchange factor Efa6 (Qu *et al*., 2019). We targeted this peptide to dendritic spines by combining it with the LifeAct peptide, which has a high affinity for filamentous actin (f-actin) (Riedl *et al*., 2008), which is concentrated in dendritic spines but is relatively sparse in the dendritic shaft. Moreover, it is well documented that MT invasion of spines is critically dependent on actin filaments in the neck of spines (Merriam *et al*., 2013; Schatzle *et al*., 2018). Thus, the depolymerizing peptide (MTED) is positioned on the f-actin that MTs require to enter spines. To control the timing of expression to mature neurons this construct is under the control of a tet-responsive promoter. We show here that expression of the LifeAct-MTED construct for only 16 hours robustly inhibits MT invasion of dendritic spines, while preserving typical MT dynamics in the dendritic shaft. Because this construct is expressed in a small percentage of neurons, it does not affect MT dynamics in presynaptic axons or synapse associated glial cells. Thus, transfection with the LifeAct-MTED construct is a much more targeted methodology for inhibiting MT dynamics than application of pharmacological compounds and can be used for studies focused on the interaction of MTs and actin filaments.

## Results

### Development of a methodology for localized disruption of microtubule dynamics

Using pharmacological compounds, such as paclitaxel or nocodazole, to inhibit MT polymerization into dendritic spines is a non-specific approach that introduces global MT disruption and a myriad of off-target effects. To address and mitigate confounding variables, we have developed a plasmid construct to specifically inhibit MT polymerization into dendritic spines, while preserving native dynamics within the dendritic shaft (**Figure 1A**). The primary component of this construct is a 20-amino-acid peptide referred to as the MT elimination domain (MTED) from the protein Efa6, which has been shown to potently depolymerize MTs (Qu *et al*., 2019). A scrambled version of the MTED peptide serves as a control in a separate plasmid **(Figure 1A)**. To selectively target the MTED or scrambled peptide to dendritic spines, we combined it with LifeAct, a 17-amino-acid peptide that readily localizes to f-actin (Riedl *et al*., 2008), which is highly enriched in the head and neck regions of spines. The fluorescent protein mScarlet (Bindels *et al*., 2017) was incorporated between the LifeAct and MTED or scrambled peptide to allow for construct visualization. To control the temporal expression of the constructs we utilized the Tet-On system, in which a doxycycline-inducible tet promoter is used in conjunction with a constitutively active hPGK promoter driving expression of the rTTA gene (David Root, RRID: Addgene_41393). The LifeAct-mScarlet-MTED or the LifeAct-mScarlet-scramble fusion protein sequence was inserted after the tet promoter, but before the hPGK promoter **(Figure 1A)**.

**Figure 1.**
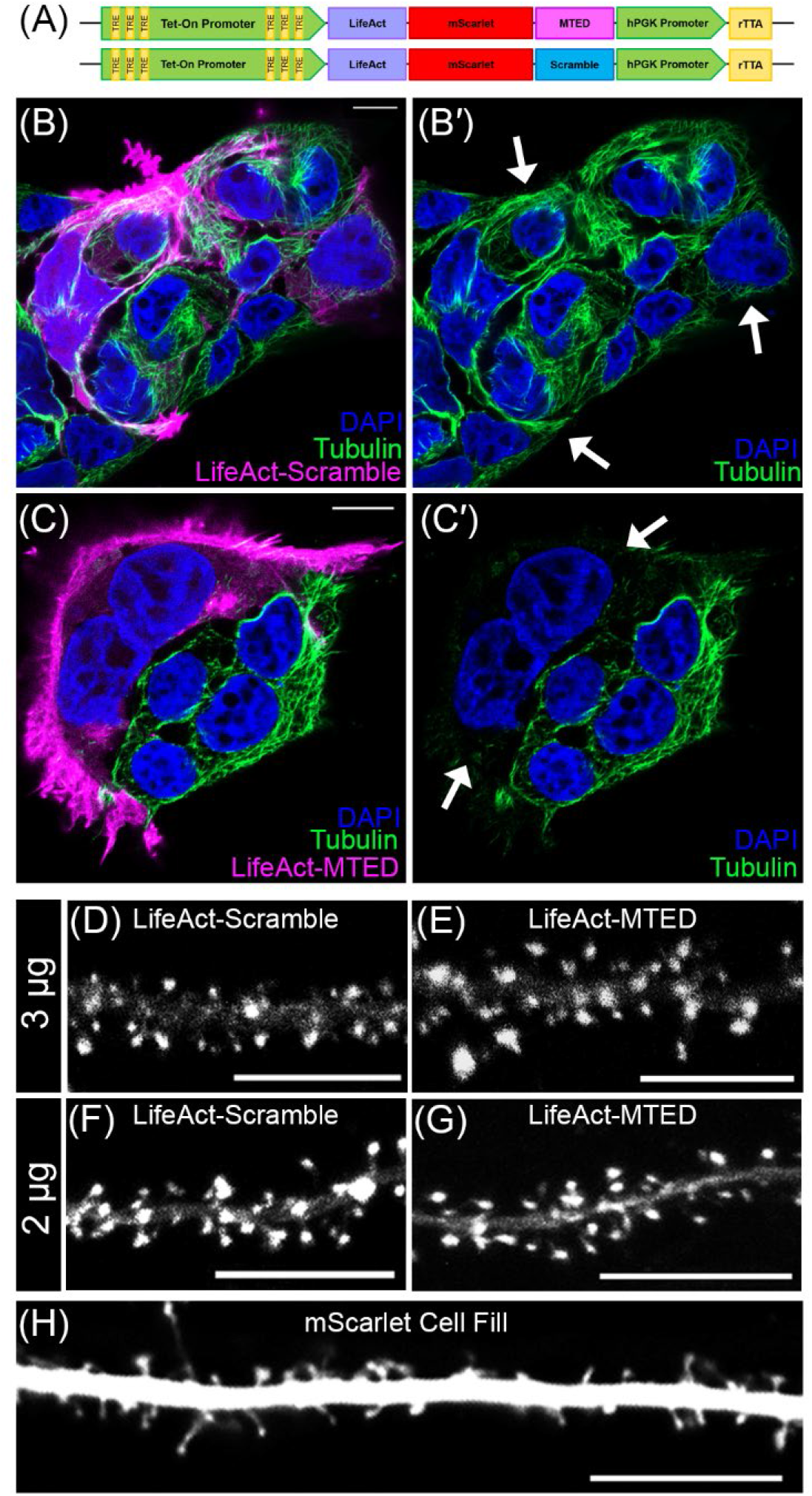
Design of the microtubule elimination domain (MTED) plasmid construct. (A) Schematic of MTED and scramble plasmid constructs containing Tet-On doxycycline inducible promoter (green) with tet-response elements (yellow), LifeAct (purple)-mScarlet (red)-MTED/scramble (magenta/blue) fusion protein sequence, separate hPGK promoter (green) and constitutively active rTTa gene sequence (yellow). (B-C’) Confocal images of fixed HEK293T cells transfected with the LifeAct-scramble (B-B’) or LifeAct-MTED (C-C’) plasmid construct and immunostained for beta-tubulin (green) with cell nuclei stained blue (DAPI). The leftmost panels (B & C) show transfected cells in magenta while the corresponding panels on the right (B’ & C’) show the same image, without the magenta overlay. Arrows in B’ and C’ point to transfected cells that contain microtubules (B’) or lack microtubules (C’). (D-H) Confocal images of living primary hippocampal neurons (DIV 21-25) transfected with 3 μg LifeAct-scramble (D), 3 μg LifeAct-MTED (E), 2 μg LifeAct-scramble (F), 2 μg LifeAct-MTED (G) or mScarlet fluorescent cell fill (H). All scale bars are 10μm.

For initial validation of the LifeAct-mScarlet-MTED and LifeAct-mScarlet-scramble plasmid constructs, which we will henceforth refer to as LifeAct-MTED or LifeAct-scramble, we utilized immortalized HEK293T cells, due to their rapid and robust culturing when compared to primary neuronal cultures. HEK293T cells were transfected with either LifeAct-MTED or LifeAct-scramble, then fixed and stained for tubulin and cell nuclei (**Figures 1B-1C′**). Tubulin content within cells transfected with LifeAct-scramble appears similar to the tubulin content of adjacent, untransfected cells (depicted by white arrows) (**Figures 1B & B′**). Conversely, cells transfected with LifeAct-MTED have a visible reduction of tubulin staining when compared to nearby untransfected cells (**Figures 1C & C′**), illustrating the MT-depolymerizing power of the MTED peptide. It should be noted that most of the MTs were depolymerized throughout the transfected cell in Figures 1C & C’. This non-specific, extensive MT depolymerization is likely due to high expression levels of the constructs in the HEK293T cells, resulting in their localization to f-actin, but with excess LifeAct-MTED dispersed throughout the cytoplasm.

While the LifeAct-MTED and LifeAct-scramble constructs appeared to be localizing to f-actin appropriately in non-neuronal cells, it was important to assess their localization in neuronal cultures. Hippocampal neurons were transfected with either LifeAct-MTED or LifeAct-scramble at two different plasmid concentrations (2 or 3 μg), allowed to mature in culture (21-25 DIV), and induced to express overnight (16 hours). We chose low concentrations of plasmid and short expression times to minimize mislocalization and disruption of MT dynamics throughout the neuron. Both concentrations of the LifeAct-scramble (**Figure 1D, F**) and LifeAct-MTED (**Figure 1E, G**) show distinct localization to dendritic spines, when compared to cells transfected with a fluorescent cell fill (**Figure 1H**). These results suggest that the plasmid constructs are localizing as expected in mature hippocampal neurons following short-term expression.

### Effect of LifeAct-MTED in mature hippocampal neurons

Once we confirmed that the LifeAct-MTED and LifeAct-scramble constructs selectively localize to dendritic spines (**Figure 1D-G**), we next investigated whether the LifeAct-MTED could effectively reduce rates of MT polymerization into spines. This was done using neurons co-transfected with a fluorescent MT end binding protein, EB3-mNeon, and varying concentrations (2 or 3 μg) of either LifeAct-MTED or LifeAct-scramble. Fluorescent EB3 binds to polymerizing ends of MTs, allowing us to visualize MT “comets” moving within the dendritic shaft and into dendritic spines in time-lapse (**Figure 2A**). Maximum projection of the collected time series shows in a single frame where MTs were polymerizing within the dendrite (**Figure 2B**). MT invasion rates were assessed by the percentage of spines invaded (invaded spines/total number of spines) and invasion frequency (total invasions/number of invaded spines) as we have done previously (Hu *et al*., 2008; Hu *et al*., 2011; Merriam *et al*., 2011; Merriam *et al*., 2013; McVicker *et al*., 2016). There is a significant reduction in the percentage of spines invaded in neurons transfected with both the higher (**Figure 2C**) and lower concentrations of LifeAct-MTED (**Figure 2E**), compared to LifeAct-scramble controls. This suggests that the MTED effectively and specifically reduces the percentage of dendritic spines being targeted for invasion by polymerizing MTs. Additionally, there is a significant reduction in invasion frequency in neurons transfected with both the higher (**Figure 2D**) and lower concentration of LifeAct-MTED (**Figure 2F**). This illustrates that there is not only a reduction in the percentage of spines being targeted for invasion, but the total number of invasions as well. Thus, it appears that the LifeAct-MTED plasmid construct is affecting invading MTs as intended and restricting MTs from polymerizing into spines.

**Figure 2.**
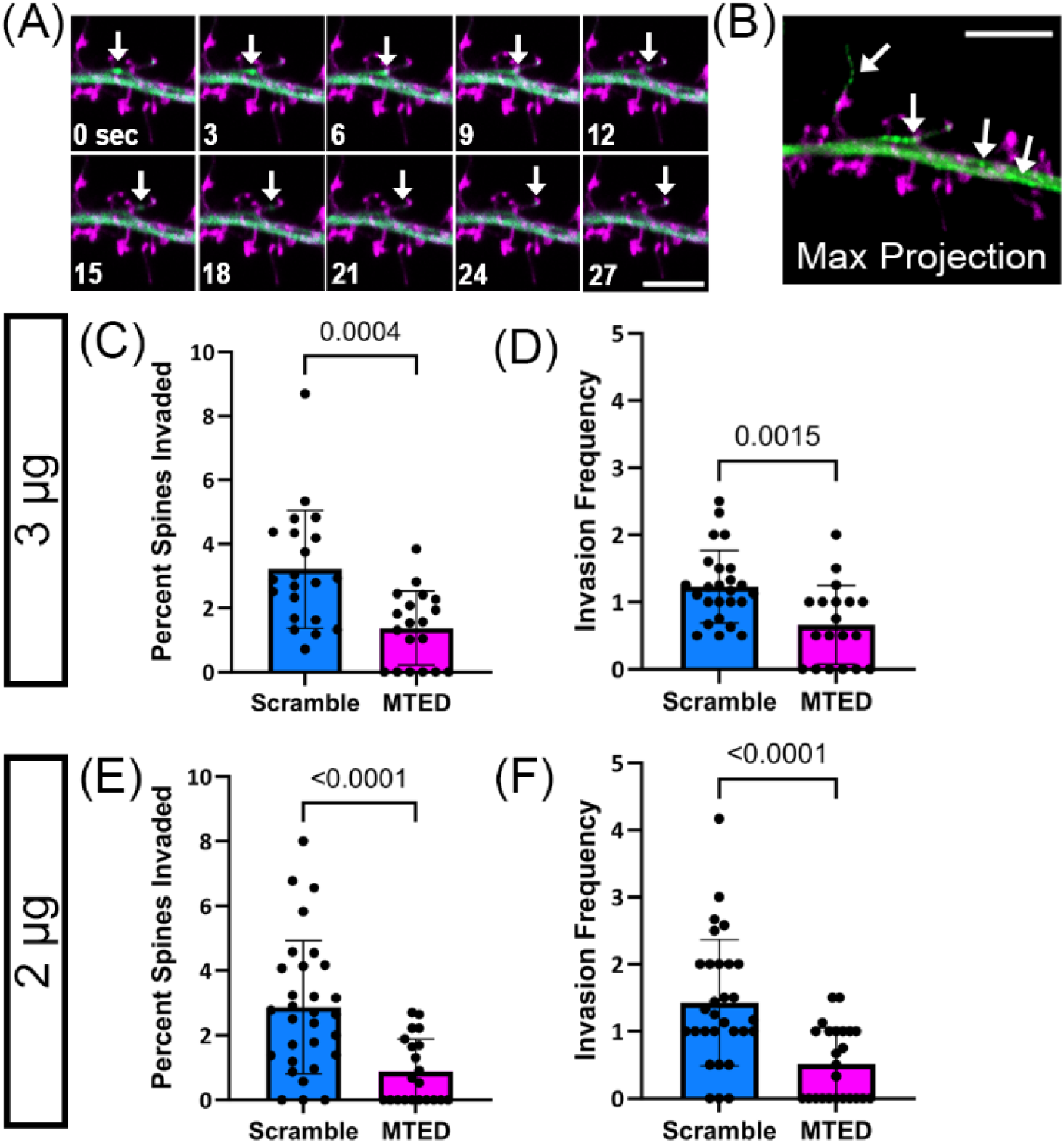
Reduction of microtubule invasions into dendritic spines within LifeAct-MTED transfected hippocampal neurons. (A) Confocal time-series of living mature hippocampal neuron (DIV 23) taken at 3-second intervals. The neuron was transfected with LifeAct-scramble (magenta) and mNeon EB3 (green) plasmid constructs. Invading EB3 comet is shown with a white arrow in each frame. (B) Corresponding maximum intensity projection of the time-series shown in A. Traces of an invading EB3 comet, as well as EB3 comets within the dendritic shaft, are shown in green and are identified by white arrows. Dendritic spines of the transfected neuron are shown in magenta. Scale bars are 5μm (A-B). (C-F) Bar graphs show mean +/-SD and black dots are individual dendritic segments. (C) Quantification of percent of dendritic spines invaded (number of invaded dendritic spines/total number of dendritic spines within field of view) for neurons transfected with 3 μg of LifeAct-scramble or LifeAct-MTED plasmid (n=21 scramble, n=19 MTED from 5 separate biological replicates for both C and D). (D) Quantification of invasion frequency (total number of invasions throughout the course of a time-series containing 100 frames/invaded dendritic spines within field of view) for neurons transfected with 3 μg of LifeAct-scramble or LifeAct-MTED plasmid. (E) Quantification of percent of dendritic spines invaded for neurons transfected with 2 μg of LifeAct-scramble or LifeAct-MTED (n=30 scramble, n=21 MTED from 5 separate biological replicates for both E and F). (F) Quantification of invasion frequency for neurons transfected with 2 μg of LifeAct-scramble or LifeAct-MTED plasmid. P values in (C-F) shown above bars are calculated with two-tailed Student’s t-test or Mann-Whitney depending on normality of data distribution.

### Examination of potential off-target effects of LifeAct-MTED in mature hippocampal neurons

Although LifeAct-MTED markedly inhibits MT polymerization into dendritic spines (**Fig 2C-F**), it remains critical to determine whether the construct has any off-target effects within dendrites. Thus, we examined MT dynamics within the dendritic shaft of transfected neurons to assess whether the typical MT dynamicity was being perturbed in the presence of LifeAct-MTED and LifeAct-scramble. Kymographs, graphical representations of EB3 comet spatial position over time, were created to visualize motion of EB3 comets within dendritic shafts of neurons (**Figure 3A**). Mature hippocampal neurons were again co-transfected with EB3-mNeon and either 2 or 3 μg of LifeAct-MTED or LifeAct-scramble. Each EB3 comet, representative of a polymerizing MT, can be visualized as a line in the kymograph, with the slope of a line representing EB3 comet velocity. The (x,y) coordinates of the beginning (x_1_,y_1_ in **Figure 3A**) and end of a line (x_2_,y_2_ in **Figure 3A**) represent the movement of EB3 comets over time. Such positions can therefore be used to calculate the distance traveled by an individual EB3 comet (x_2_-x_1_) as well as how long the comet was visualized or the “lifetime” of a comet (y_2_-y_1_) within a dendritic shaft segment.

**Figure 3.**
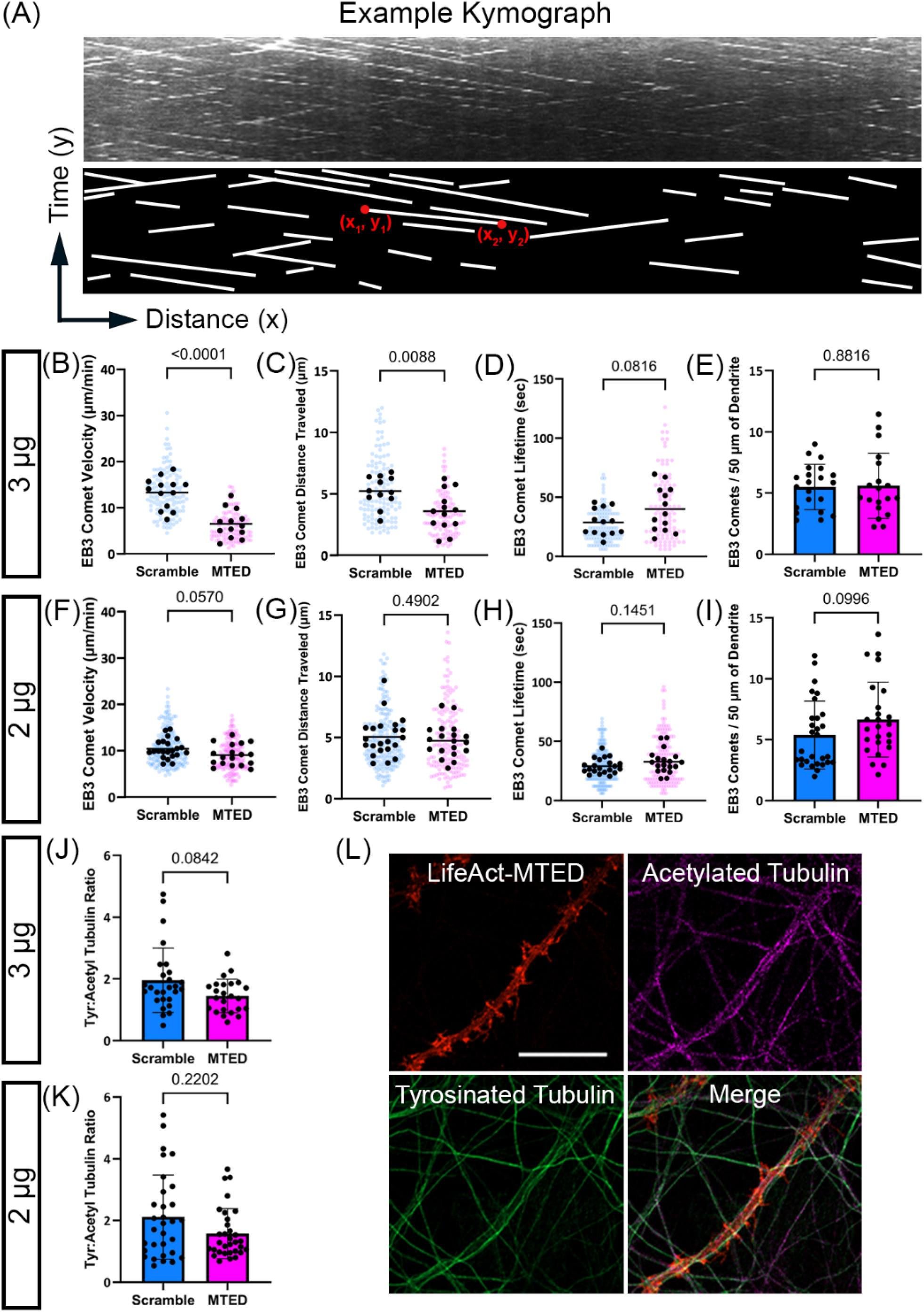
Limited off-target effects following transfection of LifeAct-MTED in mature hippocampal neurons. (A) Representative kymograph, a graphical representation of EB3 comet spatial position over time, with corresponding illustration below for simplified visualization. Each white line is representative of a single, moving EB3 comet. Coordinates of the beginning (x_1_,y_1_) and end (x_2_,y_2_) of an example line are used for calculation of distance traveled (difference in x value) and how long the comet was visualized, also referred to as the comet “lifetime” (difference in y value). (B-D) Scatter plots displaying EB3 comet velocity (B), EB3 comet distance traveled (C), and EB3 comet lifetime (D) obtained from neurons transfected with 3 μg of LifeAct-MTED or LifeAct-scramble. Mean is shown by a black bar. Each black dot is representative of the average measurement per neuron ((B) n=13 scramble, n=13 MTED; (C) n=12 scramble, n=13 MTED; (D) n=13 scramble, n=13 MTED from 4 separate biological replicates), while the blue or magenta dots in the background represent individual EB3 comet measurements ((B) n=125 scramble, n=108 MTED; (C) n=121 scramble, n=105 MTED; (D) n=118 scramble, n=107 MTED). (E) Quantification of EB3 comet abundance (number of comets per 50 μm of dendrite) within the dendritic shaft of neurons transfected with 3 μg of LifeAct-MTED or LifeAct-scramble. Bar graph shows mean +/- SD and black dots are individual dendritic segments (n=21 scramble, n=19 MTED from 5 separate biological replicates). (F-H) Scatter plots displaying EB3 comet velocity (F), EB3 comet distance traveled (G), and EB3 comet lifetime (H) obtained from neurons transfected with 2 μg of LifeAct-MTED or LifeAct-scramble. Mean is shown by a black bar. Each black dot is representative of the average measurement per neuron ((F) n=22 scramble, n=18 MTED; (G) n=22 scramble, n=18 MTED; (H) n=21 scramble, n=18 MTED from 5 separate biological replicates), while the blue or magenta dots in the background represent individual EB3 comet measurements ((F) n=195 scramble, n=164 MTED; (G) n=192 scramble, n=165 MTED; (H) n=187 scramble, n=161 MTED). (I) Quantification of EB3 comet abundance within the dendritic shaft of neurons transfected with 2 μg of LifeAct-MTED or LifeAct-scramble. Bar graph shows mean +/- SD and black dots are individual dendritic segments (n=29 scramble, n=25 MTED from 5 separate biological replicates). (J-K) Ratio of tyrosinated tubulin:acetylated tubulin within the dendritic shaft of neurons transfected with either 3 μg (J) (n=27 scramble, n=24 MTED) or 2 μg (K) (n=31 scramble, n=32 MTED) of LifeAct-MTED or LifeAct-scramble from 3 separate biological replicates. Bar graph shows mean +/- SD and black dots are individual dendritic segments. (L) Representative confocal images of fixed, hippocampal neurons transfected with the Life-Act MTED (red) and stained for acetylated tubulin (magenta) and tyrosinated tubulin (green). Such images were used for quantification of tyrosinated tubulin:acetylated tubulin ratios (J & K). P values in (B-K) shown above bars are calculated with two-tailed Student’s t-test or Mann-Whitney depending on normality of data distribution.

We observed a significant slowing of MT polymerization following transfection with the higher concentration (3 μg) of LifeAct-MTED compared to the LifeAct-scramble control (**Figure 3B**). This decrease in velocity was negated when neurons were transfected with a lower concentration (2 μg) of plasmid (**Figure 3F**), suggesting that there may be expression-level-dependent off-target effects. Thus, the LifeAct-MTED construct must be titrated to avoid disruption of MT polymerization within the dendritic shaft. It is possible this disruption of MT velocity in the dendrite is due, in part, to the presence of low, but not negligible, levels of f-actin within the dendritic shaft **(Figure 1D-G)**. To assess the mechanism by which EB3 comet velocity was decreasing with higher concentrations of LifeAct-MTED, we measured both the distance and lifetime of each polymerizing MT. Interestingly, in neurons transfected with 3 μg of LifeAct-MTED there is a significant reduction in the distance traveled by EB3 comets (**Figure 3C**). EB3 comet lifetime, however, is not significantly different compared to LifeAct-scramble control, although it does trend longer (**Figure 3D**). Thus, in neurons transfected with 3 μg of LifeAct-MTED, MTs polymerize for the same amount of time but over shorter distances, resulting in decreased comet velocity in the dendritic shaft. Together, these data suggest that MT dynamics can be significantly altered in the dendritic shaft at higher concentrations of LifeAct-MTED expression.

To decrease these off-target effects detected with 3 μg of plasmid, we transfected neurons with a lower concentration (2 μg) of LifeAct-MTED or LifeAct-scramble. We discovered with 2 μg of plasmid there is no significant difference in EB3 comet velocity **(Figure 3F)**, distance traveled **(Figure 3G**), or comet lifetime **(Figure 3H)** between LifeAct-MTED and LifeAct-scramble. To examine whether the reduction in MT invasion rates **(Figure 2C, D)** resulted from a reduction in the number of polymerizing MTs, we also measured comet abundance as a proxy for the number of polymerizing MTs in the dendrite shaft, but found no significant difference between LifeAct-MTED and LifeAct-scramble control **(Figure 3E, I)**. For additional rigor, we also compared the populations of post-translationally modified tubulin. Tyrosinated tubulin is traditionally interpreted as being incorporated into short-lived, dynamic MTs while acetylated tubulin is associated with long-lived, stable MTs (Tas *et al*., 2017). We fixed and stained for tyrosinated and acetylated tubulin in neurons transfected with either 3 or 2 μg of LifeAct-MTED (**Figure 3L**) or LifeAct-scramble to assess whether we were preferentially disrupting one specific population of MTs. However, we found no significant difference between the tyrosinated:acetylated tubulin ratio in neurons transfected with either concentration of LifeAct-MTED when compared with LifeAct-scramble controls **(Figure 3J, K)**. Taken together, these data suggest that transfection with a low concentration of LifeAct-MTED is sufficient for markedly inhibiting MT polymerization into dendritic spines, without disrupting MT dynamics and stability in the dendrite shaft.

### Assessment of correlation between various effects of LifeAct-MTED

The results of our study show that transfecting hippocampal neurons with low (2 μg) levels of the LifeAct-MTED plasmid specifically inhibits MT polymerization into dendritic spines, without affecting MT dynamics and stability in the dendrite shaft **(Figure 3F-I)**. However, expressing slightly more LifeAct-MTED plasmid (3 μg) is sufficient to significantly decrease MT polymerization velocity by decreasing the distance MTs polymerize in the dendritic shaft **(Figure 3B, C)**. One would expect that expression of either 2 or 3 μg of plasmid would result in a distribution of expression levels in individual neurons. Therefore, we measured expression levels of LifeAct-MTED or LifeAct-scramble in individual neurons and constructed scatter plots, fitted with linear regression lines against the MT metrics in spines and dendrite shafts that we measured above. We found a weak negative correlation in both the percentage of spines invaded and the invasion frequency plotted against fluorescence intensity of dendrites for the LifeAct-MTED transfected neurons compared to a weak positive correlation for the LifeAct-scramble transfected neurons **(Figure 4A-D)**. However, the slopes of the linear regression lines are not significantly different from zero, with p-values ranging from 0.0773 to 0.7578. Moreover, for neurons expressing the LifeAct-MTED plasmid we found a weak negative correlation between fluorescence intensity of dendrites and comet velocity **(Figure 4E)**, comet abundance **(Figure 4G)** and tyrosinated:acetylated tubulin ratios **(Figure 4I)**. However, neurons transfected with LifeAct-scramble have a significant (p=0.0453) negative correlation between comet velocity and fluorescence intensity **(Figure 4F)**, while having a significant (p=0.0129) positive correlation between comet abundance and fluorescence intensity **(Figure 4H)**. Nevertheless, it should be noted that the fluorescent intensity values for the LifeAct-scramble construct include average fluorescence intensity values (0-6000) much higher than for the LifeAct-MTED construct (<2000). This is because neurons expressing high levels of LifeAct-MTED (>2000) do not have discernable EB3 comets that could be used to quantify MT dynamics. Thus, at higher concentrations LifeAct-MTED is likely present in the cytoplasm and negatively affects all MT polymerization events.

**Figure 4.**
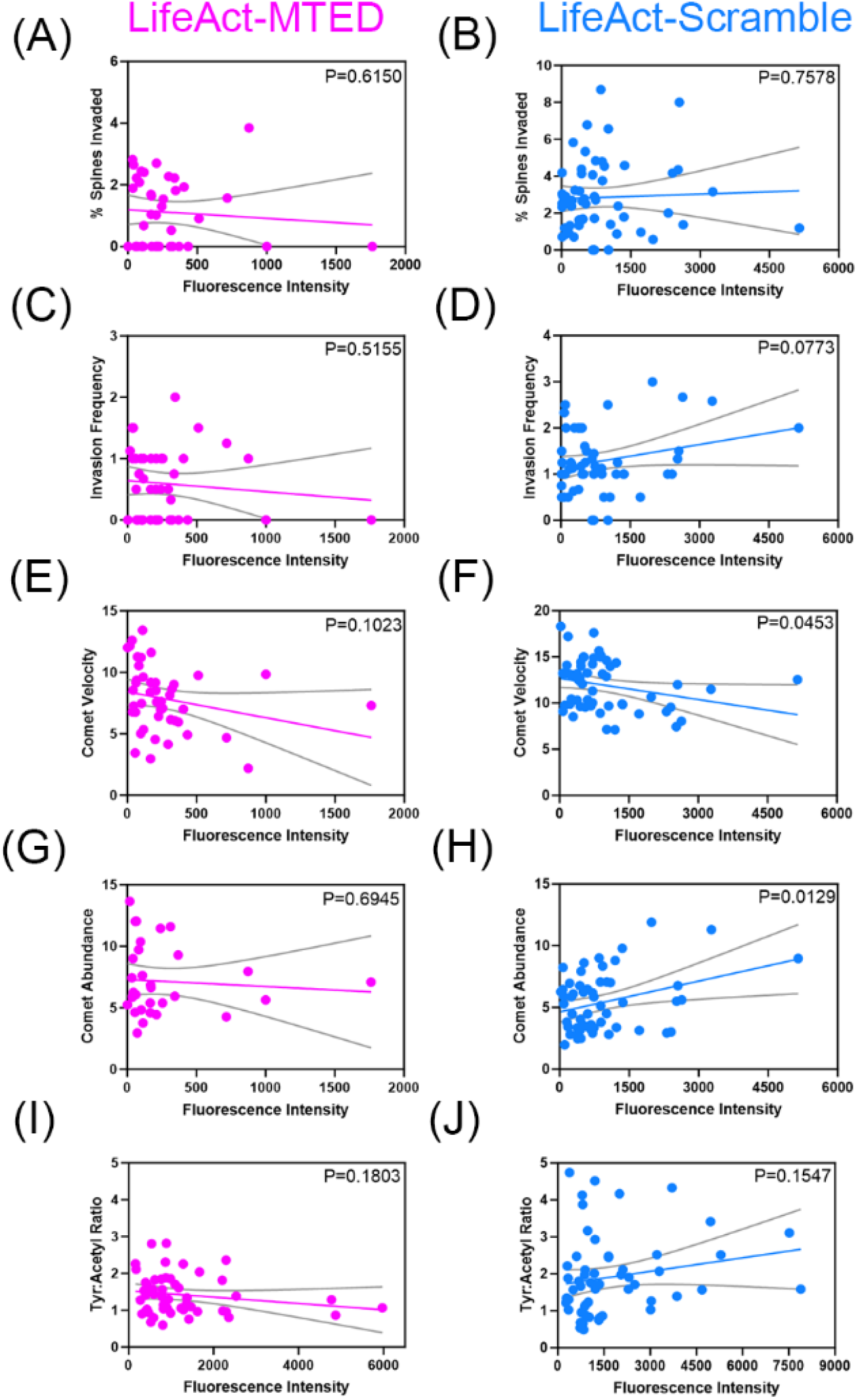
Correlation between LifeAct-MTED/scramble expression levels and microtubule dynamics. (A-J) Scatter plots of expression levels of LifeAct-MTED (A, C, E, G, I) or LifeAct-scramble (B, D, F, H, J) within hippocampal neurons, as measured by fluorescent intensity of the cell dendrite. Linear regression lines (colored) and 95% confidence intervals (gray) were plotted to compare percent spines invaded (A-B), invasion frequency (C-D), EB3 comet velocity (E-F), EB3 comet abundance (G-H) and tyrosinated tubulin:acetylated tubulin ratios (I-J) to the fluorescent intensity values. The slope of each linear regression line was analyzed for whether it significantly differed from zero, with the p-value being displayed in the upper right corner of each graph. N =40 (A), n =57 (B), n =42 (C), n =54 (D), n =44 (E), n =57 (F), n =31 (G), n =56 (H), n =52 (I), n = 57 (J) from 16 separate biological replicates.

With EB3 comet velocity appearing to be the most sensitive to variations in LifeAct-MTED expression, we sought to assess whether the magnitude of comet slowing was correlated with metrics of dendritic spine invasion. Thus, we constructed scatter plots of average EB3 comet velocity in neurons transfected with either LifeAct-MTED or LifeAct-scramble and fit linear regression lines against the percentage of spines invaded (**Figure 5A, B**) and invasion frequency (**Figure 5C, D**). We observed a slight, negative correlation of both the percentage of spines invaded and invasion frequency for LifeAct-MTED transfected neurons (**Figure 5A, C**), while there was a slight, positive correlation of both LifeAct-scramble transfected neurons (**Figure 5B, D**). While the slopes of the linear regression lines did not significantly differ from zero, we believe this importantly shows that the reduction in MT invasion rates observed with the LifeAct-MTED construct (**Figure 2C-F**) is not dependent on a reduction in EB3 comet velocity. Additionally, this weak correlation between comet velocity and measurements of MT polymerization into spines demonstrates a range of usability for our construct in that expression levels can be titrated, abolishing off-target effects, without reducing the desired effect on invasion rates.

**Figure 5.**
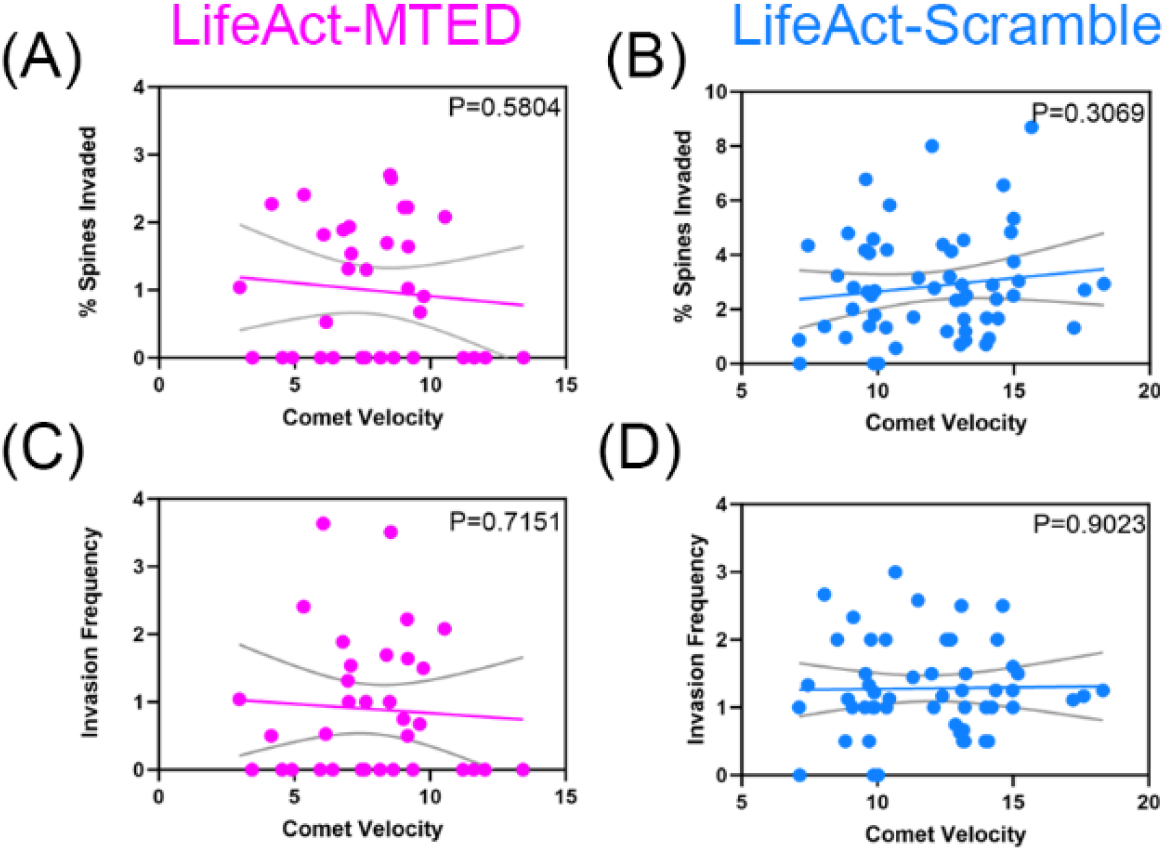
Lack of correlation between EB3 comet velocity and microtubule invasion of dendritic spines. (A-D) Scatter plots of average EB3 comet velocity within hippocampal neurons transfected with LifeAct-MTED (A, C) or LifeAct-scramble (B, D). Linear regression lines (colored) and 95% confidence intervals (gray) were plotted against metrics measuring likelihood of MT polymerization into dendritic spines, including percent spines invaded (A-B) and invasion frequency (C-D). The slope of each linear regression line was analyzed for whether it significantly differed from zero, with the p-value being displayed in the upper right corner of each graph. N =34 (A), n =57 (B), n =34 (C), n =54 (D) from 11 separate biological replicates.

## Discussion

Despite previous assumptions that MTs are largely stable polymers in mature neurons, it is increasingly appreciated that they remain dynamic throughout the life of the cell (Gu *et al*., 2008; Hu *et al*., 2008; Jaworski *et al*., 2009). Moreover, our group, as well as others, have observed MTs transiently polymerizing into, or invading, dendritic spines of hippocampal and cortical neurons (Gu *et al*., 2008; Hu *et al*., 2008; Jaworski *et al*., 2009; Hu *et al*., 2011; Kapitein *et al*., 2011; Merriam *et al*., 2011; Merriam *et al*., 2013; McVicker *et al*., 2016; Schatzle *et al*., 2018). This phenomenon appears to be highly correlated with mechanisms of learning and memory, such as LTP and LTD, in which disruption of MT dynamicity impairs learning and memory as assessed by electrophysiology or behavioral paradigms (Shumyatsky *et al*., 2005; Jaworski *et al*., 2009; Barnes *et al*., 2010; Fanara *et al*., 2010; Uchida *et al*., 2014). Such studies, however, rely on pharmacological disruption of MT dynamics. General, bath application of drugs makes it difficult to discern the origin of observed outcomes, for it effects MTs in both presynaptic axonal compartments and post-synaptic dendritic compartments, as well as any glia present within the culture. Thus, drug treatment induces many confounding variables that may contribute to, or be responsible for, the impairment of MT polymerization into dendritic spines.

To overcome the limitations associated with generalized bath application of pharmacological compounds such as paclitaxel or nocodazole, we have developed a novel plasmid, termed LifeAct-MTED, containing the discrete MT elimination domain (MTED) of the Efa6 protein (**Figure 6**). A scrambled version of the MTED is used as a control in a separate plasmid construct termed LifeAct-scramble. The MTED binds directly to tubulin, preventing polymerization, and is effective at abolishing exploratory MTs through mechanisms that remain to be fully elucidated (Qu *et al*., 2019). The MTED, coupled with LifeAct, a small peptide that readily associates with f-actin, allows for specific localization of the MTED to f-actin rich dendritic spine heads and necks (Merriam *et al*., 2013; Schatzle *et al*., 2018). LifeAct-MTED effectively inhibits not only the percentage of dendritic spines being targeted for invasion by polymerizing MTs, but also the total number of invasions occurring across the dendritic field (**Figure 2**). In addition to decreasing the percent of spines targeted and the invasion frequency of MTs polymerizing into dendritic spines, LifeAct-MTED appears to have limited off-target effects on MT dynamics within the dendritic shaft. Of the examined metrics, only EB3 comet velocity was significantly different within neurons transfected with 3 μg of LifeAct-MTED when compared to LifeAct-scramble controls. Additionally, this effect was negated upon reducing the plasmid concentration being used to transfect the neurons from 3 μg to 2 μg.

**Figure 6.**
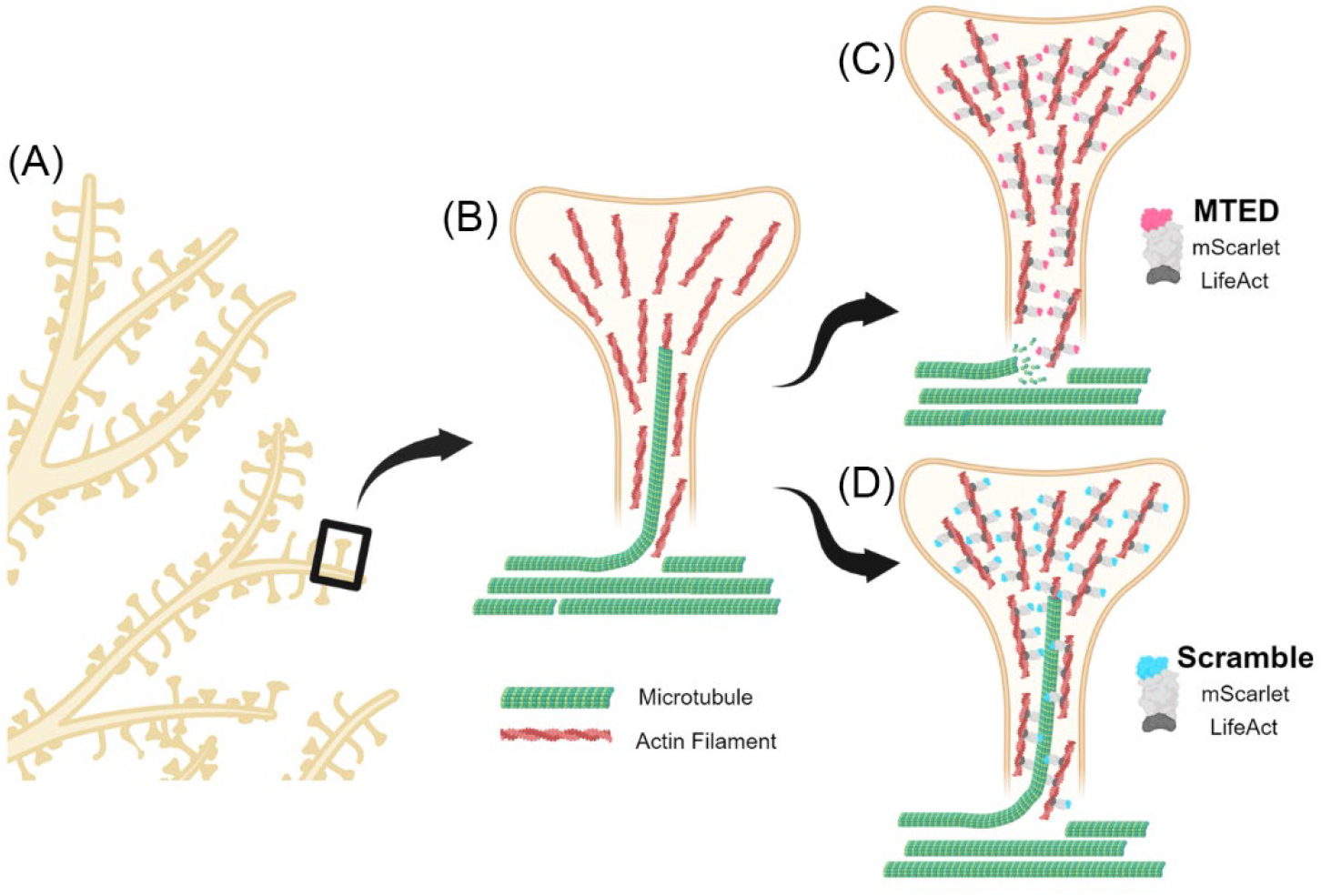
Schematic of LifeAct-MTED/scramble effects on MT invasions of dendritic spines. (A) Section of a representative dendritic arbor with one spine boxed. (B) Representative dendritic spine (yellow) rich with filamentous actin (red filaments) being invaded by a polymerizing MT (green). Other non-invading microtubules are shown in the dendrite shaft (not shown). (C) Dendritic spine (yellow) of neuron transfected with LifeAct-MTED (pink and gray fusion protein) that has localized to actin filaments (red) in the spine head and neck. LifeAct-MTED is shown depolymerizing a MT (green) prior to its entry into the dendritic spine. (D) Dendritic spine (yellow) of neuron transfected with LifeAct-scramble (blue and gray fusion protein) that is appropriately localizing to actin filaments (red) but does not affect the likelihood of a MT (green) directly polymerizing into the dendritic spine. Created with BioRender.com.

The need to titrate expression levels was largely anticipated due to the apparent potency of the MTED as well as the low, but non-negligible levels of f-actin present within dendritic shafts. Additionally, it has been observed that the MTED can be found within the cytosol when high expression levels result in saturation of the target to which it is localized (Qu *et al*., 2019). It is intriguing, however, that plasmid expression levels, measured by fluorescent intensity of transfected neurons, are not highly correlated with metrics assessed in dendritic spines and shafts (**Figure 4**). We believe this illustrates the flexibility of our tool in that there is a working range of expression that can be used to induce the desired effect on invading MTs. Nonetheless, the slowing of MT polymerization observed within neurons transfected with 3 μg of LifeAct-MTED remains an important consideration when applying this tool to other lines of research. Nevertheless, we found that the velocity of MT polymerization in dendrites was not associated with the percentage of spines invaded by MTs or in the MT invasion frequency for both LifeAct-MTED and LifeAct-scramble **(Figure 5)**. This result suggests that even though the LifeAct-MTED can decrease MT polymerization velocity at higher concentrations, this decrease is not correlated with a decrease in MT polymerization into dendritic spines. Another interesting observation was that cells transfected with LifeAct-MTED were consistently at lower levels of fluorescence in comparison to those transfected with LifeAct-scramble. We believe this was due to the difficultly of assessing MT dynamics within neurons highly expressing LifeAct-MTED, in which we could no longer visualize fluorescent EB3 comets, suggesting the LifeAct-MTED construct was appreciatively depleting MT polymerization throughout the dendrite.

Taken together, these experiments serve to illustrate that the LifeAct-MTED tool is a specific and effective means for inhibiting transiently polymerizing MTs from entering dendritic spines **(Figure 6)**. Thus, implementation of this construct could shed light on off-target effects of various MT-stabilizing and de-stabilizing drugs by comparing global disruption of MT dynamics to focal disruption only into dendritic spines. Future studies could also apply this methodology to investigate of the role of MT invasion in maintenance and/or stabilization of dendritic spine morphology. Additionally, the LifeAct-MTED construct should allow for the determination of the postsynaptic components of dendritic spines that are dependent on MT polymerization into dendritic spines. Importantly, the use of this construct is not limited to neurons and should prove useful to dissect MT/f-actin interactions in any type of cell.

## Materials and Methods

### Plasmid design and construction

MTED and scramble plasmids were assembled using standard molecular cloning techniques including PCR amplification, restriction digest and Gibson assembly. From 5’ to 3’ the constructs consist of a Tet-On doxycycline inducible promoter (a gift from David Root (Addgene plasmid # 41393 ; http://n2t.net/addgene:41393 ; RRID:Addgene_41393), a Kozak consensus sequence and a LifeAct – mScarlet (Bindels *et al*., 2017)– MTED/scramble fusion protein sequence followed by a separate hPGK promoter and a constitutively produced rTTA gene. MTED (gcgccgcgctttgaagcgtatatgatgaccggcgatctgattctgaacctgagccgcacc; APRFEAYMMTGDLILNLSRT) and scramble (atgattaccgcgccgcgcgaatttgattatctgaacctgcgcgcgggcctgagcatgacc; MITAPREFDYLNLRAGLSMT) sequences (Qu *et al*., 2019) were ordered from IDT as single-stranded ultramers with overhangs complementary to the destination vector. These, along with PCR amplified mScarlet, were inserted into Not1 and Mlu1 digested backbone by Gibson assembly reaction to create an intermediate plasmid. LifeAct peptide sequence was ordered from IDT as an ultramer with complementary overhangs and was inserted via Gibson assembly into the Ale1 and Not1 digested intermediate vector. Sequences were confirmed with Sanger sequencing (QuintaraBio). Complete maps may be found on Addgene for each respective plasmid, or are available by reasonable request.

### HEK293T cell culture, transfection, and immunostaining

HEK293T cells (Sigma, 12022001) were maintained at 37°C, 5% CO_2_ in standard HEK media comprised of DMEM High Glucose (Gibco), sodium pyruvate (Gibco) and 10% FBS. Cells were passaged at ∼80% confluency and plated onto 0.1% polyethylenimine (PEI) (Sigma) polymer-coated glass coverslips. HEK cells were transfected 24 hours after plating with Lipofectamine 3000 (ThermoFisher) transfection reagent according to the manufacturer provided protocol. Tet-On plasmid constructs were induced with 1 μg/mL doxycycline and allowed to express overnight. HEK cells were then fixed with 4% paraformaldehyde-KREB-sucrose (PKS), blocked in 10% bovine serum albumin (BSA) overnight at 4°C and incubated with primary antibody overnight at 4°C followed by secondary antibody with DAPI (1:250) overnight at 4°C. Antibodies used were mouse alpha tubulin (1:500, Sigma T9026) and goat anti-mouse AlexaFluor 488 (1:500, ThermoFisher A11029). Glass coverslips were mounted onto frosted microscope slides with Fluoromount-G (SouthernBiotech).

### Primary neuron cell culture and transfection

Primary hippocampal neurons were prepared from Sprague Dawley rats (Envigo) at embryonic day 18.5 (E18.5). Rat hippocampi were dissected and trypsinized. Dissociated neurons were resuspended in nucleofector solution (Mirus) and transfected using an Amaxa/Lonza Nucleofector II. Transfected neurons were plated at a density of 5 × 10^4^ neurons per cm^2^ on 0.1% PEI-coated glass bottom dishes (35 mm) with 14 mm micro-wells. Neurons were plated with plating media (PM; neurobasal media with 5% defined fetal bovine serum (dFBS), B27 supplement, 2 mM Glutamax, 0.3% glucose and 37.5 mM NaCl) for 2 hours at 5.0% CO2 and 37 °C after which the chambers were flooded with 2 mL of serum-free media (PM with no added dFBS). Neurons used within experiments ranged from DIV 21-25. All procedures were approved by the University of Wisconsin Committee on Animal Care and were in accordance with the NIH guidelines.

### Tyrosinated:acetylated tubulin immunocytochemistry

Primary hippocampal neurons were allowed to develop to DIV 21-25, and then induced with 1 μg/mL doxycycline for overnight expression. After approximately 16 hours, neurons were fixed with 4% PKS for 20 minutes (Dent and Meiri, 1992). Blocking solution of 10% BSA was applied and allowed to incubate overnight at 4°C. Primary antibody was incubated at 4°C, as was secondary antibody. Antibodies used were rat tyrosinated alpha-tubulin (1:1000, Millipore MAB1864-I), mouse acetylated alpha-tubulin (1:2000, ThermoFisher 32-2700), goat anti-rat AlexaFluor 488 (1:500, ThermoFisher A11006) and donkey anti-mouse AlexaFluor 647 (1:200, ThermoFisher A32787).

### Confocal imaging

Images were acquired on a Zeiss LSM 800 confocal microscope with a 63x/1.4NA Plan Apochromat oil objective. For spine invasion and comet velocity/abundance data, quick successive time-lapse images were acquired in a single channel (EB3-mNeon) at a rate of 1 frame/3 sec for a duration of 100 frames. During live, time-lapse microscopy, neurons were kept at 37°C in a warmed chamber enclosing the microscope and with a glass ring sealed with silicone grease and a glass coverslip to maintain appropriate CO_2_ levels. Fixed cell imaging for collection of acetylated:tyrosinated tubulin data utilized z-stacks that were acquired in red, green, and far-red channels with slice increments of 0.24μm.

### Image analysis

Images were processed using ImageJ software (NIH). Spine invasions were defined as distinct EB3 puncta or “comets” at least two times brighter than the background fluorescence of the dendritic shaft moving into a dendritic spine and persisting there for at least two frames. The percentage of spines invaded was determined by dividing the number of invaded spines by the total number of spines in the dendritic field. Invasion frequency was defined by the total number of invasions divided by the number of spines invaded. EB3 comet velocities were calculated by the slope of the line produced by the kymograph tool in ImageJ and did not include stationary or paused events. EB3 comet distance traveled was derived from the difference in x values of the coordinates at the beginning and end of the kymograph line and EB3 comet lifetime was derived from the difference in y values. Fluorescence intensity measurements were collected from confocal microscopy images. The average background fluorescence was subtracted from the entire image. Normalized fluorescence was measured in five regions of interest along each dendritic field. Fluorescence intensities were averaged between three dendritic sections per neuron.

### Graphing and statistical analysis

All statistical tests and graphing were performed in Prism 8 (GraphPad). Outliers were identified by the ROUT method in which Q = 1% and subsequently excluded. Data were tested for normality using the Kolmorgorov-Smirnov test of normality. If data were normal, then a two-tailed *t-*test was performed. If data were not normal, then a Mann-Whitney U test was performed. P values for scatter plots were determined by simple linear regression analysis in Prism. Data with *P* values less than 0.05 were considered statistically significant. Complete data are available upon request.

## Abbreviations

BSA: bovine serum albumin
f-actin: filamentous actin
FBS: fetal bovine serum
LTP: long-term potentiation
LTD: long-term depression
MT: microtubule
MTED: microtubule elimination domain
PKS: paraformaldehyde/Krebs/Sucrose
Tet: tetracycline

## Author Contributions

EH, MM and ED contributed to the conception and design of the study. EH and HM executed the experiments. RT and GD assisted with analysis of confocal microscopy images and provided laboratory support. EH and ED wrote the manuscript with input from HM. All authors read and approved of the final submitted version. ED supervised all aspects of the work.

## Acknowledgements

This work was supported by NIH grants R01-NS098372 and R01-NS115400 to ED. EH was supported, in part, by NIH training grant T32-NS105602. We thank Innes Hahn (University of York) and Andreas Prokop (University of Manchester) for providing reagents and advice at the beginning of this project.

